# Intrinsic specificity of a ‘core’ Tip60 acetyltransferase complex in *Drosophila*

**DOI:** 10.1101/2025.08.19.671066

**Authors:** Silke Krause, Marco Borsò, Marisa Müller, Axel Imhof, Zivkos Apostolou, Peter B. Becker

## Abstract

The lysine acetyltransferase Tip60 (KAT5) regulates gene expression through acetylation of histone N-terminal ‘tail’ domains. We determined the intrinsic substrate selectivity of a recombinant, 4-subunit TIP60 core module from *Drosophila melanogaster* with synthetic nucleosome arrays. We compared matched arrays of nucleosomes containing either the replication-dependent histone H2A, or the variant H2A.V (H2A.Z in mammals), a prominent substrate of Tip60.

Targeted mass spectrometry allowed to quantify acetylation of individual lysines in histones H2A, H2A.V and H4. Overall, H4 and H2A/H2A.V were equally well acetylated. The analysis comprehensively identified selected sites of acetylation, their relative acetylation levels, diacetylation patterns and revealed surprisingly different acetylation rates of individual lysines. We also applied this defined acetylation system to evaluate the effectiveness and selectivity of a TIP60 inhibitor, NU9056. Remarkably, the inhibitor shows variable effectiveness at different acetylation sites. Knowledge about the intrinsic substrate selectivity of Tip60 is a prerequisite for a mechanistic understanding of the enzyme’s mode of action.

## Introduction

The histone acetyltransferase KAT5 governs crucial aspects of eukaryotic genome functions, including the regulation of genome-wide transcription and the response of cells to DNA damage (1–4). The importance of the enzyme is illustrated by the fact that it has been conserved during evolution from the unicellular yeast *Saccharomyces cerevisiae* to humans. The KAT5 of yeast, Esa1, is the catalytic subunit of the yeast NuA4 complex (5–7). In mammals, KAT5 is also known as TIP60. The epigenetic P400 regulator combines KAT5 with an ATP-dependent histone H2A.Z variant exchange activity, which is contributed by the p400 ATPase (8).

Unlike other KAT enzymes that acetylate nucleosomes at specific sites, KAT5/TIP60 appears to have a fairly relaxed specificity, since in diverse model systems and assays it has been reported to acetylate several lysines on histone H2A (2,9), the variant histone H2A.Z (Htz in yeast: (10–14) and histone H4 (2,15–17). This relaxed specificity contrasts other KATs that belong to the same ‘MYST’-family of acetyltransferases. For example, MOF (KAT8) exclusively acetylates the H4 N-terminus.

The seemingly relaxed selectivity of KAT5/TIP60 on histone H4 and H2A N-termini, which protrude from the nucleosome particle on the same side (18), may be explained by the fact that the substrate lysines in the N-terminal tail domains of histones H4, H2A and H2A.Z are embedded in very similar local sequence motifs (Figure 1A). Nevertheless, depending on whether H2A or H4 is to be acetylated, KAT5/TIP60 must approach nucleosomes in very distinct ways. Recent structural work suggests that a 4-subunit complex in yeast consisting of Esa1, along with the subunits Yng2, Eaf6, and Epl1, first acetylates H4 and then reorients itself on the mononucleosome substrate to subsequently acetylate H2A.Z (19).

**Figure 1.**
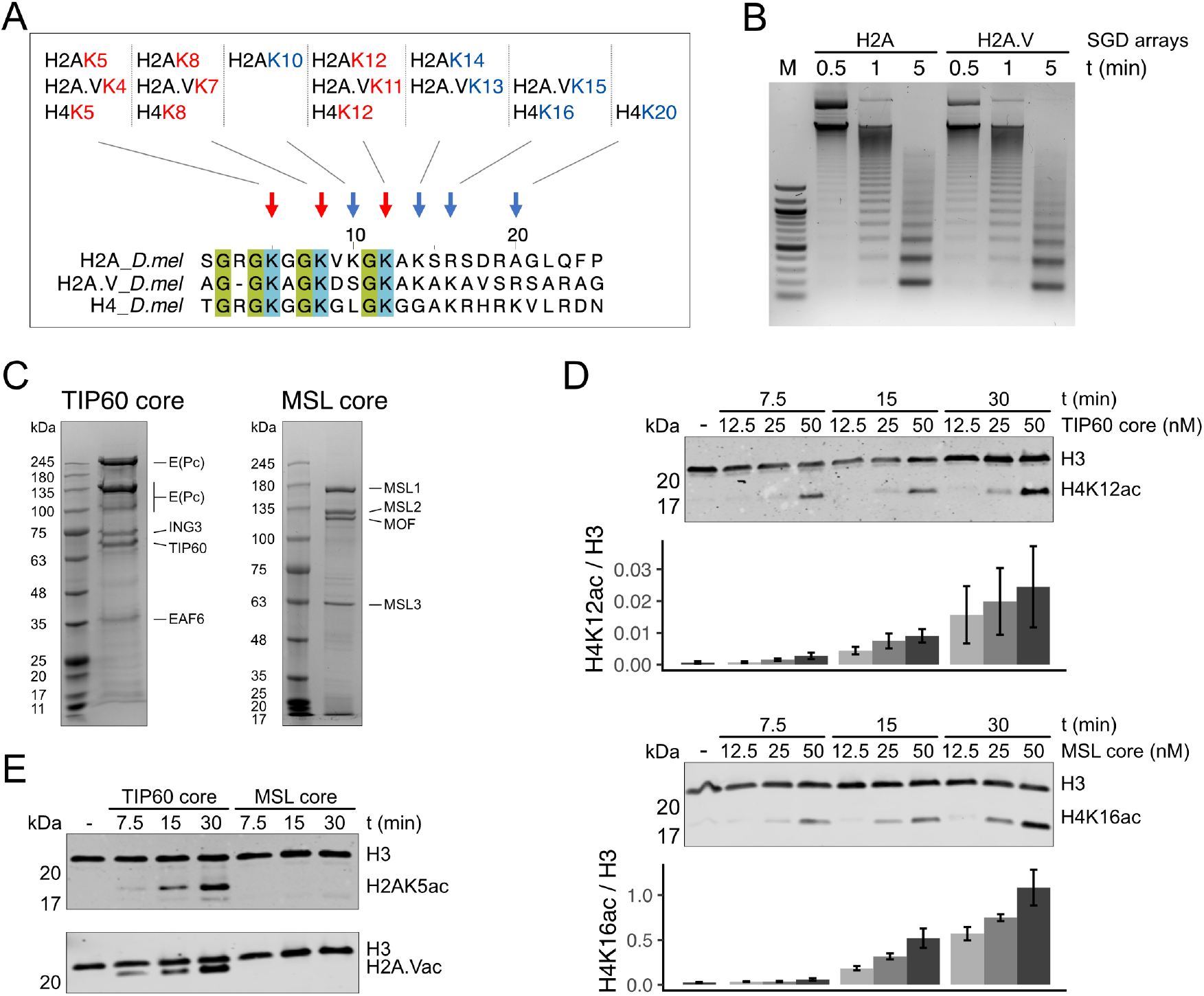
The TIP60 core complex acetylates H2A, H2A.V and H4 N-terminal histone tails in vitro. A) Graphical representation of the N-terminal histone tails H2A, H2A.V and H4. The sequences shown correspond to the N-terminal tails of the histones, with the initial methionine omitted, which is post-translationally cleaved off. Positions of lysines marked in red are found at similar positions in H2A and H4, blue residues are H2A-or H4-specific. B) Examples of purified TIP60 and MSL core complexes used in this study. C) Nucleosome arrays were assembled by salt gradient dialysis (SGD) using either H2A- or H2A.V-containing *Drosophila* histone octamers. The arrays were subjected to Micrococcal nuclease (MNase) digestion to assess nucleosome occupancy and array organization. D) Acetylation of histone H4. Nucleosome arrays were incubated with the indicated concentrations (nM) of TIP60 or MSL core complexes for specified reaction times (minutes). Acetylation of histone H4 was assessed by immunoblot using antibodies specific to H4K12ac (upper panel) and H4K16ac (lower panel). Error bars represent the standard error of the mean (SEM) from 4 biological replicates for the TIP60 core complex and 3 biological replicates for the MSL core complex. E) Nucleosome arrays containing H2A (upper panel) or H2A.V (lower panel) were incubated with 50 nM of purified TIP60 or MSL core complexes for the indicated times. The immunoblots used antibodies raised against H2AK5ac and H2A.VK4acK7ac (H2A.Vac), respectively.

In *Drosophila melanogaster*, Tip60 resides in a nucleosome remodeling complex akin to the mammalian P400 complex, which in flies is called DOMINO-A for historical reasons (16). In line with the evolutionary conservation of the N-terminal histone domains, TIP60 has been shown to acetylate H4K5 and H4K12 *in vivo* (16) and *in vitro* (15). When it comes to the other prominent substrate, the histone variant H2A.Z, *Drosophila* presents an interesting case: H2A.V, the H2A.Z-like protein in flies, contains a C-terminal sequence epitope that is phosphorylated in response to DNA damage, similar to the mammalian H2A.X. The fly H2A.V thus appears to be chimaera of mammalian H2A.Z and H2A.X, and accordingly combines the functions of the two mammalian histone variants in promoter architecture and DNA damage signaling (20). It is possible that the selectivity of acetylation in cells is modulated by chromatin context and circumstances.

We explored the intrinsic substrate selectivity of a *Drosophila* Tip60 ‘core’ complex. As is the case in yeast (7), the individual Tip60 enzyme can acetylate lysines on peptides representing the N-termini of histones H2A, H2A.Z, H3 and H4 with relaxed selectivity, but it cannot modify histones organized as nucleosomes (5,7). We previously showed that a 3-subunit assembly of *Drosophila* Tip60 orthologous to the yeast ‘piccolo’ complex (5,7) acetylates H4K12 on oligonucleosome substrates (15). In our current experiments we co-express the Eaf6 subunit to reconstitute a 4-subunit *Drosophila* Tip60 ‘core’ complex [according to the nomenclature of Xu *et al*. (21)], consisting of E(Pc), Ing3, Tip60 and Eaf6. Since relaxed selectivity may be observed on low-complexity substrates (such as histone peptides or free histones), we reconstitute nucleosome arrays that homogenously contain either canonical H2A or the variant H2A.V. Because antibody-based experiments are compromised by the non-equivalent avidity and specificity of the antibodies, we employ targeted mass spectrometry (15,22) to quantitatively determine the extent of acetylation of individual lysines in the N-termini of H2A, H2A.V and H4.

This set-up allows to address several open questions related to the substrate selectivity in nucleosome arrays. The variant H2A.V differs from canonical H2A in the extent of the acidic patch, a crucial interaction site on nucleosomes. Does this translate into changes in acetylation? Does the presence of H2A or H2A.V affect the efficiency or kinetics of H4 acetylation? Is there a hierarchy, either kinetically or quantitatively, between H2A.V and H4 on the same nucleosome substrate, as suggested by Wang and colleagues (19)? At which lysines are H2A and H2A.V acetylated and do the two variants differ in this respect? Is oligo-acetylation detectable, as has been described in a similar experimental set-up with the Mof-containing MSL complex (15)?

The analysis of Tip60 effects on cells would profit very much from the availability of a specific inhibitor. Taking advantage of the defined *in vitro* acetylation system we evaluate the effectiveness and specificity of two published KAT5 inhibitors.

## Methods

### Cloning, protein expression and purification of the TIP60 core complex

The TIP60 core complex was cloned using the biGBac technology (23). The cDNAs of full-length *Drosophila melanogaster* E(Pc), Ing3, Eaf6 and Tip60 fused to an N-terminal TwinStrep tag were combined in one pBIG1 vector. Bacmids were generated in DH10 Multibac cells. The proteins were co-expressed in SF21 cells, infected with 1/1000 (V/V) baculovirus lysate for 72 h at 26°C. Cells were collected by centrifugation, washed once with PBS buffer, flash frozen in liquid nitrogen and stored at −70°C.

The TIP60 core complex was purified from 10^8^ baculovirus-infected SF21 cells. Cells were resuspended in 10 ml Lysis buffer [25 mM Hepes pH 7.6, 300 mM NaCl, 5% glycerol, 0.1% NP40, 3 mM MgCl_2_, Complete EDTA-free protease inhibitor (Roche)] and supplemented with 2.5 *µ*l RNase A (10 *µ*g/*µ*l) and 1 *µ*l Benzonase (Merck). After incubation for 15 min on ice, the suspension was sonicated with the Branson sonifier (50 sec total,10 sec on, 20 sec off, 20% amplitude). Cell debris was pelleted by centrifugation 30,000g for 30 min. The supernatant was filtered through a Minisart filter of 1.2 *µ*m pore size. The clear supernatant was loaded onto a 0.5 ml Strep-Tactin XT4 gravity flow column, equilibrated with Wash buffer (25 mM Hepes pH 7.6, 250 mM NaCl, 5% glycerol, 3 mM MgCl_2_). The column was washed twice with 10 column volumes (CV) Wash buffer and eluted with 4 × 0.5 CV BXT-Buffer (100 mM Tris pH 8.0,150 mM NaCl,1 mM EDTA, 3 mM MgCl_2_, 50 mM biotin). The quality of the protein preparation was assessed by SDS-PAGE and Coomassie staining. The concentration was determined in Image Lab 6.0 (Bio-Rad) using BSA standards (ThermoFisher) as reference.

The core MSL complex was expressed and purified by FLAG-tag chromatography as before (24).

### Histone expression and octamer assembly

We employed histones from *Drosophila melanogaster*. The H2A.V expression plasmid was a gift from Reinhard Kalb and Jürg Müller (MPI Martinsried, Germany). The expression of histones and octamer assembly was as described (25) with the following modifications: due to the lower pI, histone H2A.V were purified in buffer Sau-0 instead of Sau-200. Histones H3 and H4 were kind gifts from Dr. C. Regnard (BMC Molecular Biology, Martinsried), prepared by the purification of inclusion bodies. For octamer reconstitution, ratios of the corresponding histones were titrated to reach final ratios of H2A:H2B:H3:H4 1.2:1.2:1:1. Appropriate quantities were pooled, lyophilized and resuspended in unfolding buffer to final concentrations of 4.7 mg/ml for H2A and H2B, and 4.0 mg/ml for H3 and H4, respectively. The histones were dialyzed against refolding buffer at 4°C overnight. Octamers were purified by size exclusion chromatography in refolding buffer on a Hiload 16/600 Superdex 200 column (Sigma).

### Assembly of nucleosome arrays by salt gradient dialysis

Nucleosome arrays were assembled by salt gradient dialysis on a pUC18 plasmid comprising 25 repeats of a 197 bp long Widom-601 nucleosome positioning sequence. Ten *µ*g plasmid DNA was mixed with octamer in a 1.1:1 mass ratio in a High Salt buffer (100 *µ*l, 10 mM Tris-HCl pH 7.6, 2 M NaCl, 1 mM EDTA, 0.05% IGEPAL CA630, 14.3 mM β-mercaptoethanol) and transferred to dialysis cups (Slide-A-Lyzer; MWCO 3500, Thermo Fisher). The cups were floated in a beaker containing 300 ml High Salt buffer but with 0.1% 2-mercaptoethanol). The NaCl concentration in the beaker was decreased constantly at room temperature (RT) overnight by pumping 3 L of Low Salt buffer (10 mM Tris-Cl pH 7.6, 50 mM NaCl, 1 mM EDTA, 0.05% Igepal CA-630, 0.01% 2-mercaptoethanol) into the beaker with a peristaltic pump (Minipulse evolution, Gilson, mode 7.6). After the Low Salt buffer had been transferred, the dialysis cup was dialysed for another 1 h at RT against fresh Low Salt Buffer. The concentration was determined based on DNA absorption at 260 nm wavelength. Chromatin was stored at 4°C and used for up to 12 weeks.

Nucleosome array quality was evaluated by Micrococcal Nuclease (MNase) digestion. 500 ng nucleosome arrays were mixed with 0.005 units MNase (Sigma-Aldrich) in MNase buffer (20 mM HEPES pH 7.5, 50 mM NaCl, 3 mM MgCl_2_, 2.5 mM DTT, 0.5 mM EGTA, 1.5 mM CaCl_2_) and incubated for 30 s, 60 s or 5 min at 30°C. The reaction was stopped by addition of 10 mM EDTA and 0.4% (w/v) SDS. The sample was treated with 2.5 *µ*L Proteinase K (10 mg/mL, Bioline) for 30 min at 37°C. DNA was ethanol-precipitated, centrifuged at 21,000 g at 4°C for 30 min, and washed once with 400 *µ*L 70% ethanol. The pellet was air-dried and resuspended in 15 *µ*L 10 mM Tris-HCl pH 8.0 and 3 *µ*L of Orange G loading dye (NEB). The digestion degree was analyzed on a 1.5% (w/v) agarose gel in TAE buffer, stained with Midori green (Nippon Genetics).

### Acetyltransferase (HAT) assay

TIP60 core complex was mixed with 0.8 *µ*g nucleosome arrays, 0.2 *µ*g RNase A and 10 *µ*M acetyl-CoA (Sigma-Aldrich) in HAT buffer (50 mM HEPES pH 7.9, 3 mM MgCl_2_, 50 mM KCl) at 26°C. Protein concentration (12.5–100 nM) and incubation times (5–60 min) varied between the different experiments. The reaction was stopped by heating to 95°C for 5 min. Histone acetylation was analyzed by Western Blot or by mass spectrometry.

### Western blotting

The denatured HAT reaction samples were mixed with SDS-PAGE loading buffer and boiled before loading onto a 15% (v/v) acrylamide Bis-Tris-MES gel in MES buffer (50 mM MES, 50 mM Tris, 1 mM EDTA, 0.1% SDS, 5 mM sodium metabisulfite). Proteins were separated at constant 120 V and transferred to a nitrocellulose membrane in Transfer buffer (25 mM Tris, 192 mM glycine) for 1 h, 400 mA at 4°C). Membranes were blocked in 3% (w/v) BSA in TBS for 1 h at RT and probed overnight at 4°C with antibodies specific for histone H3 (anti-H3, mouse, Abcam ab10799), histone H4 acetylation (anti-H4K12ac, rabbit, Millipore 07-595) and histone H2AZac, (rabbit, Cell Signaling 75336), respectively. Membranes were washed thrice with TBS-T and incubated with species-specific secondary antibodies (LICOR). Images were recorded with the LICOR imager.

### Sample preparation for mass spectrometry

Samples for mass spectrometry were prepared as described (15). Briefly, heat-denatured HAT assay samples were chemically acetylated using 25% (v/v) fresh acetic anhydride-D6 (Sigma-Aldrich) for 45 min at 37 °C with 500 rpm agitation. The pH was adjusted stepwise to pH 7.0 by addition of 1 M ammonium bicarbonate. Light and heavy isotope-labeled acetyl groups, which differ in mass by 3 Da, have identical chemical properties allowing for reliable MS quantification and discrimination between chemical and enzymatic acetylation. Proteins were digested with 250 ng of trypsin for 16 h at 37 °C with 500 rpm agitation. Peptides were desalted using commercially available C18 Attract SPE disks (Affinisep) which were washed with 60 μl each of (i) 100% acetonitrile, (ii) 0.1% (v/v) trifluoroacetic acid, 80% (v/v) acetonitrile in MS-grade water and (iii) 0.1% (v/v) trifluoroacetic acid in MS-grade water. At each step, liquid was passed through by centrifugation at 300xg for 3 min at RT. Trypsin digested samples were loaded to the tip, centrifuged 10min at 300xg and the flow through was loaded once again. Bound peptides were washed three times with 0.1% (v/v) trifluoroacetic acid in MS-grade water. Finally, peptides were eluted in three steps with 0.25% (v/v) trifluoroacetic acid, 80% (v / v) acetonitrile in MS-grade water and dried under vacuum in a speed vacuum at 45 °C. Samples were resuspended in 17 μl of 0.1% (v/v) trifluoroacetic acid in MS-grade water, sonicated for 5 min and stored until measured at −20°C.

### Mass spectrometry analysis of histone modifications

Samples were injected in an Ultimate 3000 RSLCnano system (Thermo) coupled to a Qexactive HF (Thermo). Peptides were separated using a 25-cm Aurora column (Ionopticks) with a 90-min gradient from 2 to 43% acetonitrile in 0.1% formic acid. The MS instrument was programmed to target several ions as described previously (15). For the analysis of H2A and H2A.V, triply charged ions of their corresponding acetylated forms were selected to be fragmented. Of note, H2A.V was chemically acetylated at the N-terminus and the corresponding peptides were selected for the targeted analysis (Supplementary Table 1). Survey full scan MS spectra (from m/z 450–1100) were acquired with resolution *R* = 30 000 at *m*/*z* 400 (AGC target of 3e6). Targeted ions were isolated with an isolation window of 1.3 m/z to a target value of 2e5 and fragmented at 27% normalized collision energy. Typical mass spectrometric conditions were: spray voltage 1.5 kV; no sheath and auxiliary gas flow; heated capillary temperature, 250 °C.

### Data analysis of MS data

Raw data from mass spectrometry was analyzed using Skyline (15) v25.1. Peak integration was performed for H4, H2A and H2A.V peptides for each of their corresponding modifications. Relative levels of each PTM were calculated from the obtained intensities using R (26), based on the formula given in (15). Formula to calculate the relative levels of H2A and H2A.V acetylated peptides are shown in Supplementary Figure 1 and 2 respectively.

## Results

### The TIP60 core complex acetylates histones H2A, H2A.V and H4 in nucleosome arrays

The previously described, somewhat promiscuous acetylation of histones H2A, H2A.Z and H4 by TIP60 may to some extent be explained by sequence similarities in the N-terminal ‘tail’ sequences (Figure 1A). The three histones display conserved lysines K5, K8 and K12, preceded by glycine residues, which may allow flexible accommodation of the substrate in the active site of the acetyltransferase. Notably, *Drosophila* H2A contains a unique K10 residue that is absent in H2A.V and H4 and not conserved in the human H2A homolog.

To investigate the intrinsic substrate selectivity of the *Drosophila* TIP60 core complex on most physiological substrates, we assembled nucleosome arrays containing either histone H2A or the H2A.Z-like variant H2A.V from recombinant histone octamers by salt gradient dialysis. Because we aimed for a quantitative comparison of acetylation, we carefully matched the nucleosome occupancy of the arrays, which is assessed by micrococcal nuclease (MNase) digestion (Figure 1B) and histone H3 immunoblotting (Figure 1E and data not shown). These arrays were used as substrates for a recombinant, affinity-purified *Drosophila* TIP60 core complex consisting of E(Pc), Ing3, Eaf6, and Tip60 (Figure 1C). We compared this TIP60 complex to the MSL core complex, composed of the acetyltransferase Mof and associated proteins Msl1, Msl2 and Msl3, known for its defined substrate selectivity towards H4K16 (15) (Figure 1C).

To establish the appropriate enzyme concentration and incubation time for the mass spectrometry experiments, we carried out a series of pilot immunoblot experiments using commercial antibodies raised against H4 ‘tail’ lysines, known to be preferred substrates for either Tip60 (H4K12) or Mof (H4K16). As expected, TIP60 core acetylated H4K12, while MSL core showed robust activity on H4K16ac (Figure 1D). Subsequent detection of histone H3 provided an internal reference. In both cases, acetylation levels increased with protein concentration and incubation time. Quantification across independent experiments confirmed the reproducibility of this activity. The H4K12ac signal (H4K12ac/H3) maximally reached 3% at highest TIP60 concentration and 30-minute incubation time. In contrast, the Mof-dependent H4K16 acetylation was about 30-fold stronger. While it is possible that these activity differences are due to intrinsic properties of the enzymes (see (15)), we cannot be sure because antibody avidities may be very different.

We next evaluated the activities of TIP60 core and MSL core to acetylate H2A and H2A.V. Immunoblot analysis with antibodies against H2AK5ac revealed that the TIP60 core complex was capable of acetylating the H2A or its variant H2A.V, detected by antibodies raised against H2AK5ac and H2A.VK4acK7ac, respectively (Figure 1E). The MSL core complex failed to acetylate H2A/H2A.V. Together, these findings demonstrate that the reconstituted *Drosophila* TIP60 core complex is competent to acetylate H2A, the H2A.V variant and H4K12 in our assay system. Whereas both TIP60 core and MSL core can acetylate the H4 tail, Tip60 stands out for its ability to modify H2A.

### Complex acetylation dynamics of TIP60 core on histone H4 and H2A substrates

To quantify lysine-specific histone acetylation by the TIP60 core complex, we employed targeted mass spectrometry (MS), thus bypassing the limitations of antibody specificity. Mindful of potential variability of enzyme and substrate preparations, our quantitative data represent the average of two biological replicates consisting of two different enzyme preparations assayed with two independent nucleosome array assemblies. H2A and H2A.V arrays were always assembled and assayed in parallel for consistency. We chose 50 nM TIP60 core complex to obtain a robust signal (Figure 1D, E) and monitored acetylation during a time course, assuring linearity of the assay.

We first examined H2A-tail acetylation on nucleosome arrays containing canonical H2A (Figure 2A). The MS-based heatmap revealed selective acetylation at lysines K5, K8, and K10, with K10 showing the highest overall modification level, followed by K8 and K5. No acetylation was detected at K12 or K14, even after extended reaction times. Interestingly, K5-K8 diacetylation was clearly detected. Due to a lack of differentiating fragment ions K5-K10 and K8-K10 diacetylations could not be distinguished by MS2, and are therefore represented as combined value (Figure 2A). These findings suggest that TIP60 core does not acetylate all accessible lysines on the H2A tail indiscriminately, but acetylates individual N-terminal lysines up to and including K10, which also contribute to di-acetylated forms.

**Figure 2.**
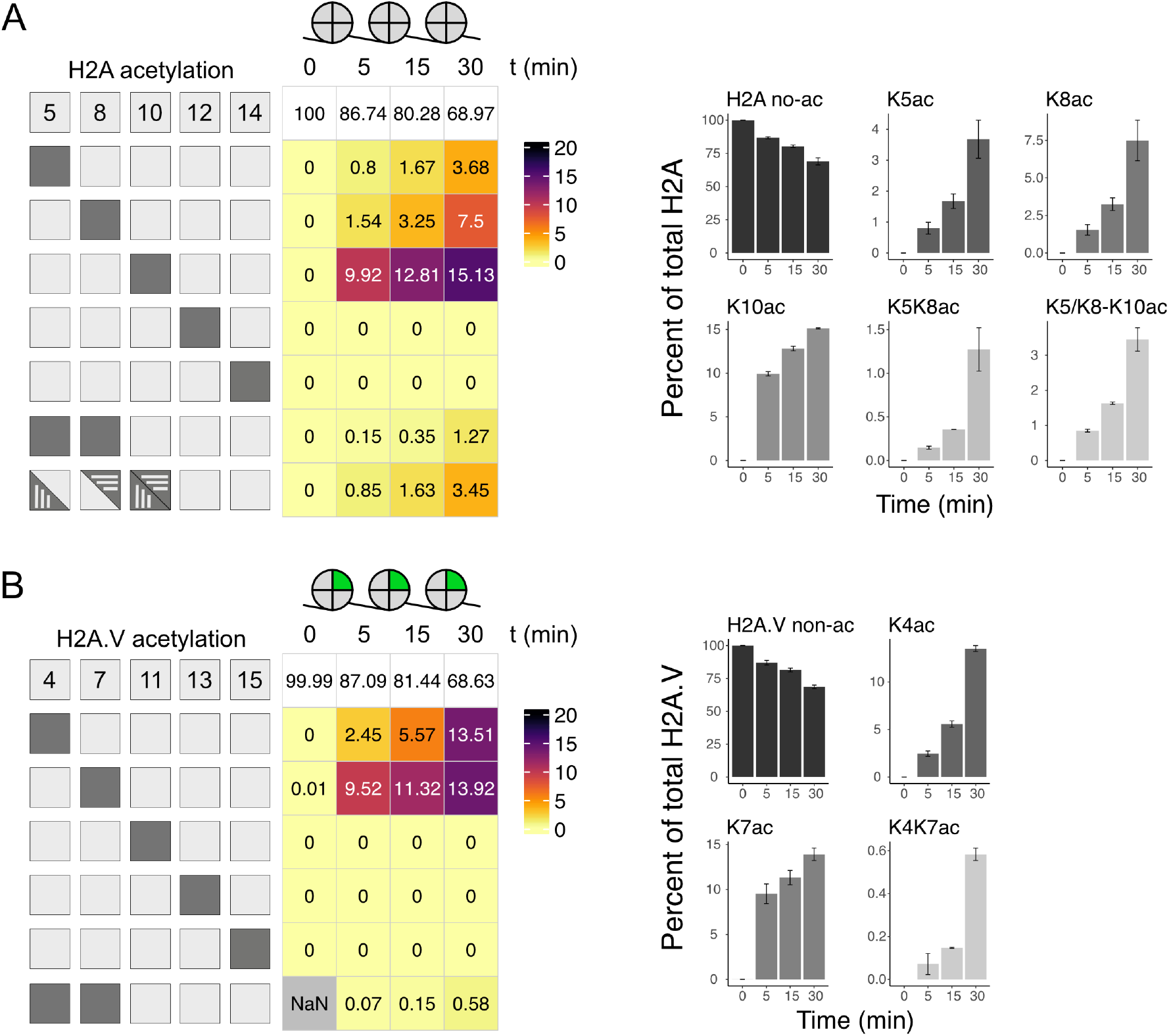
Lysine-selective acetylation of histone H2A and H2A.V by TIP60 core. A) Left panel: Heatmap showing site-selective acetylation of H2A lysines 5, 8, 10, 12, and 14 in H2A-containing nucleosome arrays by 50 nM TIP60 core complex, as quantified by targeted MS2 analysis. Light shading indicates non-acetylated residues, while dark shading indicates acetylation at the corresponding lysine position. Only acetylation combinations detectable by MS2 are shown. The heatmap represents the mean relative abundance of acetylated motifs at indicated time points of 2 biological replicates (N = 2) with 2 technical replicates each. The top row displays the mean levels of remaining non-acetylated lysines. NaN: No light or heavy fragments detected. 0: No light fragment detected, but a heavy fragment detected. Right panel: Bar plots summarizing the abundance of detected mono- and di-acetylated H2A tail forms. Note different scales of the graphs. Error bars represent values from two independent biological replicates (2 TIP60 core complexes and 2 nucleosome arrays), each analyzed with two technical replicates. B) As in (A) displaying site-specific acetylation of H2A.V tail lysines in H2A.V nucleosome arrays.

To determine how the histone variant H2A.V influences substrate selectivity, we repeated the MS analysis on nucleosome arrays reconstituted with H2A.V (Figure 2B). The TIP60 core complex acetylated K4 and K7 on the H2A.V tail. Notably, at 5 minutes, K7 was acetylated approximately four times more efficiently than K4, but by 30 minutes, both residues reached comparable levels of modification. This kinetic difference may reflect different H2A.V-specific enzyme-substrate interactions. Unexpectedly, K4-K7 diacetylation accumulated to lower levels relative to H2AK5K8 diacetylation, suggesting that monoacetylation of K4 or K7 is the preferred acetylation form of the TIP60 core complex *in vitro*.

Analysis of H4 acetylation on H2A-containing arrays revealed a clear preference for K12, which was robustly acetylated in a time-dependent manner (Figure 3A). K5 was also acetylated but remained relatively stable over the time course, while K8, a residue flanked by these two sites, was only weakly modified. K16 showed low-level but consistent time-dependent acetylation. Evidently, the TIP60 core complex acetylates lysines in the H4 N-terminal tail selectively, in contrast to the sequential pattern described earlier for Mof in the MSL core complex, which primarily targets H4K16 and acetylates more outward lysines in a processive manner (15), akin to the ‘zip’-model described by Burlingame and colleagues in butyrate-treated HeLa cells (27). Nevertheless, combinations of di- and triacetylation were much more frequent if K12ac was included, suggesting that K12 is the primary target.

**Figure 3.**
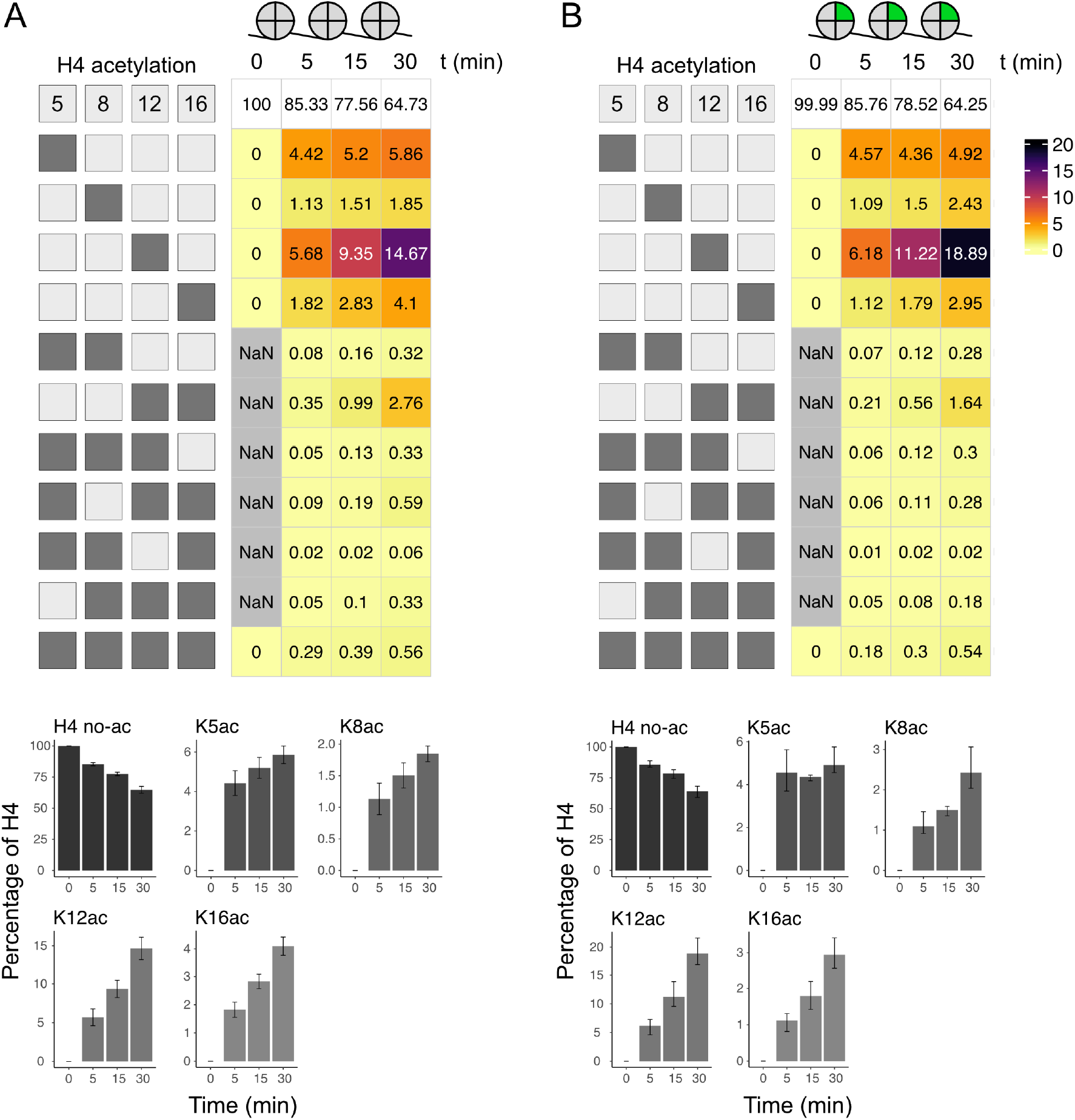
Lysine-selective acetylation of histone H4 by TIP60 core as a function of H2A variant. A) Heatmap and bar plots as in Figure 2A displaying site-specific acetylation of H4 tail lysines in H2A nucleosome arrays. B) Heatmap and bar plots as in Figure 2A displaying site-specific acetylation of H4 tail lysines in H2A.V nucleosome arrays.

The pattern of H4 acetylation in the context of H2A.V arrays was largely similar to that observed with H2A arrays (Figure 3B). H4K12 was again the primary site of modification, followed by lower levels of acetylation at K5 and K16. However, the extent of K12 acetylation was 22% higher in the H2A.V context, suggesting that the presence of H2A.V in the nucleosome may facilitate or stabilize TIP60 core binding and activity on adjacent H4 tails.

Together, these findings demonstrate that the TIP60 core complex acetylates nucleosomes with a surprising lysine selectivity. The preference for certain lysines over others, especially the poor acetylation of H4K8 and absence of H2A K12/K14 acetylation, argues for a model in which structural constraints and local sequence context strongly influence TIP60 core substrate recognition and catalytic activity. These results further suggest that H2A variant composition within chromatin can subtly modulate acetylation efficiency and specificity.

### NU9056 inhibits TIP60 core histone acetylation with lysine-specific effects

Given the central role of Tip60 in chromatin regulation and genome maintenance, the development of chemical inhibitors targeting its acetyltransferase activity has attracted considerable interest. It is difficult to conclude about direct effects of an inhibitor from studies in cells. Our defined *in vitro* system is ideally suited to evaluate the effectiveness of candidate inhibitors of the intrinsic activity of Tip60. We focused on two chemical compounds, TH1834 (28) and NU9056 (29) that had been suggested in the literature as inhibitors for the human TIP60, yet their specificity had not been unequivocally established.

In pilot experiments, we added increasing concentrations of the two compounds to *in vitro* acetylation assays with either TIP60 core or MSL core complexes (Figure 4A, B). TH1834 had no detectable effect on either complex, even at very high concentrations. In contrast, 2 *µ*M (but not 0.2 *µ*M) NU9056 completely eliminated TIP60 core-mediated acetylation of H4K12 and H2A.V. Acetylation of H4K16 by MSL core was only affected at about 100-fold higher NU9056 concentrations. These results suggest that NU9056 inhibits TIP60 much more effectively than Mof,

**Figure 4.**
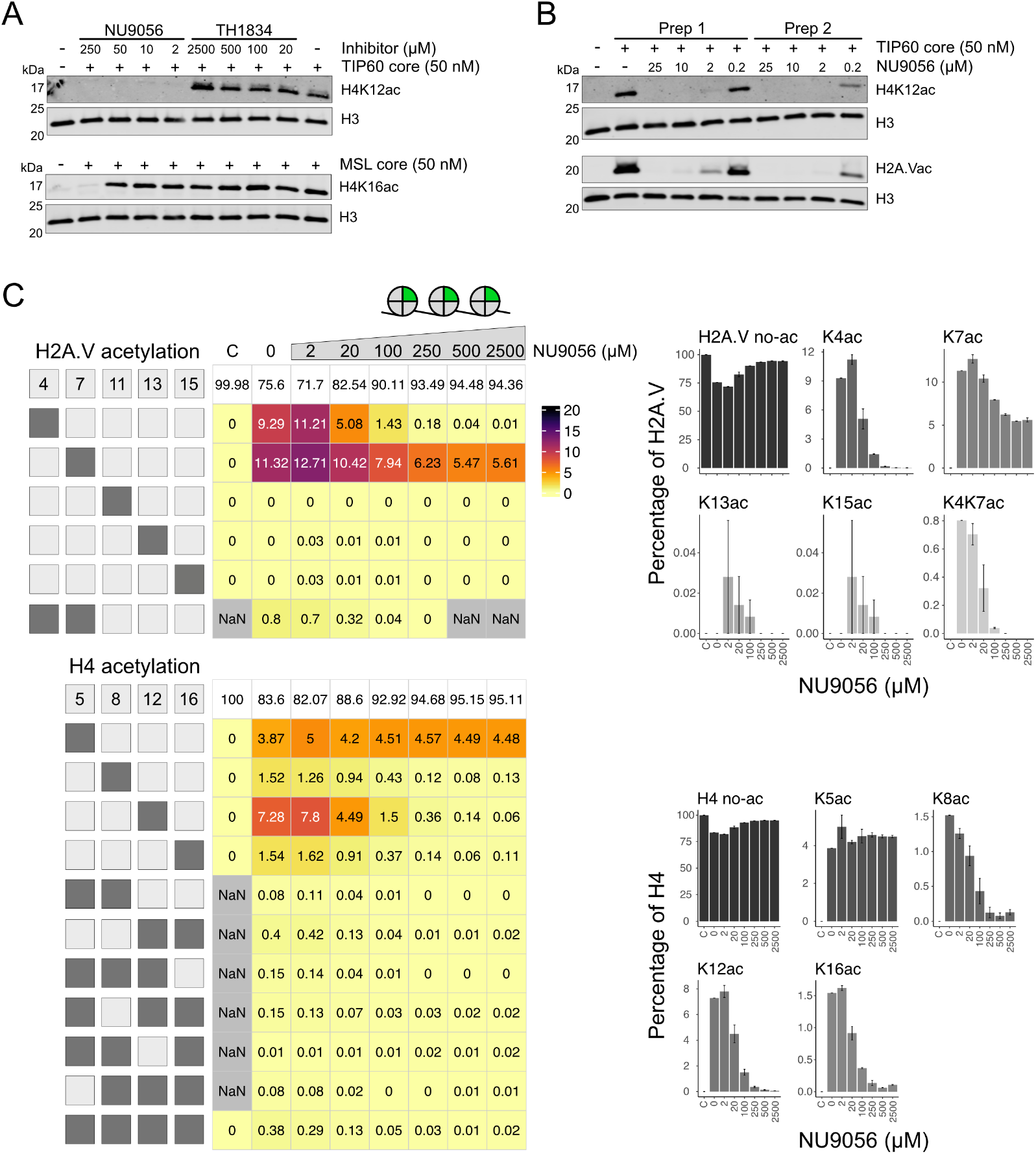
Effect of NU9056 on the TIP60 core acetyltransferase activity. A) Immunoblots representing histone acetylation assays performed in the presence of increasing concentrations of NU9056 or TH1834. Reactions were carried out on H2A.V nucleosome array substrates using 50 nM purified TIP60 or MSL core complexes for 15 minutes. Acetylation of H4K12 was assessed for the TIP60 core complex, and H4K16 acetylation was assessed for the MSL core complex using specific antibodies. B) Immunoblots showing the effect of increasing NU9056 concentrations on the acetylation activity of two different TIP60 core complex preparations on H2A.V nucleosome array substrates. Acetylation of histone H4K12 and H2A.V was assessed using site-specific antibodies. C) Heatmaps (left) and bar plots (right) as in Figure 2A showing the effect of indicated concentrations of NU9056l on lysine acetylation on the H2A.V (top) and H4 (bottom) with 50 nM TIP60 core complex. Only acetylation patterns detectable by MS2 are shown, while some multi-acetylated forms are not captured. The heatmaps represent the mean abundance of acetylated motifs at the indicated time points. The top row of each heatmap shows the mean levels of non-acetylated lysines under each condition. Two independent TIP60 core preparations were used (N = 2). Error bars represent the two replicate values.

To describe the effect of NU9056 on TIP60 core activity more quantitatively, we repeated the MS analysis on H2A.V-containing nucleosome arrays incubated with increasing concentrations of NU9056 (Figure 4C). Remarkably, while acetylation of H2A.VK4 and H4K12 were reduced more than two orders of magnitude by increasing inhibitor concentrations, the acetylation of H2A.VK7 was only modestly reduced (2-fold) and the acetylation of H4K5 was not affected at all, even at the highest inhibitor concentration. This observation resonates with the unusual acetylation kinetics at these sites we observed earlier: unlike other lysines, H4K5 and H2A.VK7 gain most of their acetylation within 5 minutes and acetylation does not (H4K5) or only moderately (H2A.VK7) increase during the remaining incubation time.

In summary, NU9056 effectively inhibits the intrinsic catalytic TIP60 core activity at low concentrations, but also impairs Mof activity at higher concentrations. Moreover, its inhibitory effect varies across different lysine residues, even on the same histone tail.

## Discussion

Protein acetylation bears potential to form an intricate regulatory network that allows cells to adapt to changing conditions (30). A subset of this modification system involves acetylation of histones to locally open chromatin to promote transcription. The relationship between acetyltransferase ‘writers’, deacetylase ‘erasure’ enzymes and proteins that specifically recognize and bind acetylated lysines is complex, which precludes easy identification of substrates for a KAT of interest through *in vivo* analyses. For example, the p300 acetyltransferase (KAT3) colocalizes with high histone H4 acetylation due to binding of a bromodomain subunit, but then acetylates the histone variant H2A.Z (31). KAT enzymes may be functionally redundant or cross-regulate each other through acetylation (3,30).

As a first step towards deciphering the acetylation network the intrinsic substrate specificities of KAT enzymes may be determined in defined *in vitro* systems. Our analysis for the acetyltransferase Tip60 (KAT5) from *Drosophila* are relevant for the mammalian system due to the high evolutionary conservation of the enzyme and its substrates.

To approximate the physiological substrate, we assembled matched nucleosome arrays bearing either H2A or H2A.V. These arrays are efficiently acetylated by the 4-subunit TIP60 core complex, which forms an autonomous KAT module that is flexibly tethered to the larger DOM-A complex through the E(pc) subunit (32,33). We employed targeted mass spectrometry to unequivocally identify not only individual acetylated lysines, but also di- and oligo-acetylated species (15,22). This approach avoids uncertainties about the selectivity of antibodies that are commonly used to identify acetylated lysines (34,35). We made several interesting observations that warrant structural and biophysical follow-up studies.

### Histone H2A acetylation

Histone H2A K5, K8 and K10 are individually acetylated by the TIP60 core complex. Acetylation at these sites as well as at K14 had been shown to be functionally redundant in yeast (14). We did not observe significant modification of K12 (K11 in H2A.V), despite the fact that this residue should be accessible (18) and bears the conserved ‘GK’ signature, which allows fitting the substrate lysine into the active site of the enzyme (21). This selectivity illustrates the requirement for a defined interaction of the TIP60 core complex with the nucleosome substrate (19,21).

We also did not score K14 acetylation. Acetylation of the homologous H2AK15 has been detected in human 293T cells and shown to be placed by a recombinant human TIP60 complex with human histones *in vitro* (9). However, this acetylation was achieved with a TIP60/EP400 complex that resembles the DOM-A complex in *Drosophila*. The presence of the nucleosome remodeling ATPases P400 may lead to a degree of nucleosome remodeling that facilitates acetylation of the H2A tail closer to the nucleosome surface.

Remarkably, we observed that the individual H2A lysines K5, K8 and K10 were acetylated with different kinetics and to different extent during the 30-minute reaction. H2A acetylation is usually detected with an antibody raised against an K5ac-bearing peptide. Ironically, K5 is the least acetylated lysine among the three. The acetylation strength follows a gradient from the most ‘inward’ K10 to the peripheral K5, reminiscent of the way MOF in the context of the MSL core module modifies the H4 N-terminus (15). Interestingly, H2AK10 is neither found in H2A.V nor in H4, suggesting a specific function yet to be discovered.

### Variant histone H2A.V acetylation

Whereas the replication-dependent H2A resides in most nucleosomes in the genome, H2A.V, the only H2A variant histone in *Drosophila*, is incorporated by exchange of H2A, independent of replication (20,36). *Drosophila* H2A.V resembles mammalian H2A.Z for most of its sequence, but carries a C-terminal epitope that is phosphorylated in the context of the DNA damage response, similar to mammalian H2A.X (20). The N-termini of H2A and H2A.V show similar ‘GK’ motifs up to K11/12, beyond which the sequences diverge. H2A.V (like H2A.Z in other species) contributes to a larger ‘acidic patch’ on the flank of nucleosomes, an important docking site for interacting proteins (37). This could be the reason why H4K12 in H2A.V-bearing nucleosomes is acetylated a little better than H4 in canonical nucleosomes.

The commercially available monoclonal antibody to detect H2A.Z acetylation was raised against a K4/K7-diacetylated human H2A.Z peptide. It can be used to detect H2A.V acetylation due to cross-reactivity. We now found that each of the two lysines is individually acetylated with similar rates, but in the presence of NU9056 acetylation of K4 is inhibited two orders of magnitude, whereas K7 acetylation is found only 2-fold reduced. These subtleties are not revealed using immunoblotting.

*Drosophila* H2A.V lacks K10, which is so avidly acetylated in H2A, but the remaining K4 and K7 are acetylated to higher degrees, as if to make up for the lack of K10. Interestingly, a comprehensive analysis of histone modification by mass spectrometry in human cells only detected acetylation of K4 and K7 despite the presence of K11 in human H2A.Z (22). The functional significance of these phenomena is unclear at present.

The acetylation reactions proceed very differently on the two histone variants: after 30 minutes H2A.VK4 is acetylated 4-fold more than the corresponding H2AK5. The low levels of diacetylated peptides reveal that K4 and K7 of H2A.V rarely appear acetylated on the same histone tail. This contrast corresponding reactions of Mof in the MSL context, where di-, tri and even tetra-acetylated species accumulate over time due to the processive action of the enzyme (15). Similarly unexplained are the different acetylation kinetics of individual lysines in close neighborhood on the same histone tail. Whereas K7 is acetylated fast and the reaction plateaus at 30 minutes, K4 acetylation starts slower and continues to increase until it reaches similar levels to K7.

### Histone H4 acetylation

The better-known substrates for Tip60 so far had been H4K12, H2AK5 and H2A.VK4/K7, but due to non-comparable antibody avidities, it was unknown whether the tails would be acetylated to similar degrees. We found this to be the case. Despite interesting differences in kinetics, lysine residues on H2A/H2A.V and H4 are acetylated to a roughly similar extent during the 30-minute reaction. H4K12 is acetylated about 22% better in the context of H2A.V, reminiscent of the early observation that H4 acetylation was higher on native H2A.Z-versus H2A-containing nucleosomes in cells (38). Structural studies aimed to decipher how the yeast piccolo NuA4 module engages the nucleosome to modify the two distinct tail substrates suggest that the complex adopts different conformations on substrate nucleosomes. Apparently, the enzyme first binds and acetylates the H4 tail and then reorients to modify the H2A.Z tail (19). The suggested hierarchy of acetylation is not obvious from our study.

We recently described the kinetics, with which Mof in the context of the MSL core complex acetylates H4 of an H2A-containing array (15). A comparison with the TIP60 core complex is instructive since both enzymes are MYST acetyltransferases and bear significant similarity in their catalytic center (19). The core MSL complex acetylates the H4 tail mainly at K16, but also modifies the more ‘outward’ lysines with decreasing efficiency. It also accumulates significant di- and oligo-acetylated forms during extended incubation. Mathematical modeling suggests that this acetylation profile arises form extensive a processive reaction starting from H4K16. According to this line of thought, the TIP60 core complex does not act in a processive mode, as it has a very pronounced selectivity for K12 and other lysines are acetylated less, independent of their position in the H4 N-terminus. The acetylation of the H4 tail illustrates the limitations of antibody-based approaches. Tip60 acetylates H4K12 about 5-fold better than H4K16. NU9056 inhibits K12 acetylation about a 100-fold and K16 acetylation only 10-fold. Using only an H4K16ac-specific antibody one might be satisfied with this effect, and while it is possible that K16 acetylation by Tip60 has functional relevance, the main effect is clearly on K12.

### Functional implications

H2A.V (and its acetylated form) is enriched in the ‘+1’ nucleosome just downstream of active promoters (30), where it may contribute to destabilizing the nucleosomal barrier in preparation for promoter escape of RNA polymerase II (36). In contrast, H2A is distributed throughout the genome. In the yeast model, the exchange of H2A for H2A.V was strongly stimulated by Tip60-dependent acetylation of H2A and H4 (39). The functional redundancy between H2A and H4 in this assay may explain the seemingly relaxed selectivity of Tip60. In the context of a DOM-A complex targeted to a promoter, Tip60-dependent acetylation of H2A and H4 would generate binding sites for the bromodomain subunit Brd8. The stabilized binding of the complex might promote the incorporation of H2A.V and, once the exchange has been achieved, continue to acetylate H2A.V to establish the quality of the ‘+1’ nucleosome. In yeast, such a scenario is executed by functional interaction between the NuA4 and SWR1 complexes (12,36,40). In this context it will be interesting to explore whether the presence of H3.3, also a hallmark of the ‘+1’ nucleosome, affects the substrate selectivity.

In addition to this promoter-centric scenario, Tip60 may function through alternative routes. We recently showed that Tip60 has numerous non-histone substrates in nuclei. Many of the acetylated proteins have known roles in gene expression, cell cycle progression and energy provision (30).

### Evaluation of candidate Tip60 inhibitors

RNAi-mediated depletion of Tip60 leads cell proliferation stop at various points in the cell cycle (30). A more acute loss of function would be desirable. Although some KAT5 inhibitors have been reported, the literature does not allow to conclude about Tip60/KAT5 selectivity (41,42). In any case, our defined *in vitro* system is ideally suited to evaluate the effectiveness of candidate inhibitors.

We focused on two chemical compounds that had been suggested in the literature as inhibitors for Tip60, yet their specificity had not been unequivocally established. Addition of TH1834 (28) to our reactions did not have any effect on H4K12 acetylation. This compound had been initially characterized in the chicken system, which may explain why it does not work on the *Drosophila* enzyme.

The second compound that is advertised as Tip60 inhibitor, NU9056, inhibits cell proliferation and is widely used to treat cancer. The compound inhibited the activity of recombinant Tip60 to incorporate tritiated acetate into free histones 50-fold better than recombinant GCN5, p300 or PCAF. If LNCaP cells were treated with NU9056, levels of H4K8ac, H4K16ac as well as H3K14ac, were decreased, as seen by immunoblotting (29).

We observed that NU9056 inhibited acetylation of H4K12 by the TIP60 core module at 50-fold lower concentrations than the MSL core complex. In a pilot experiment, we monitored the inhibition of Tip60 using mass spectrometry. The fact that acetylation of individual lysines are inhibited with different kinetics argues against a mechanism involving competition with acetyl-CoA, but for an allosteric distortion of the catalytic center. The explanation of these unusual kinetics of acetylation and enzyme inhibition will require further structural and biophysical studies in the context of NU9056.

## Concluding remarks

Our study has answered several open questions about the intrinsic selectivity with which the TIP60 core complex acetylates chromatin. Explaining the rich phenomenology of lysine-specific acetylation kinetics will require follow-up biophysical land structural studies.

## Acknowledgments

We would like to acknowledge Dmytro Hlushchenko for his effort in testing the effects of the Tip60 inhibitors during the early phase of the project.

## Author contributions

S.K. (Data curation, Investigation,). M.B. (Data curation, Formal analysis, Investigation,). M.M. (Validation). A.I. (Validation). Z.A. (Conceptualization, Data interpretation, Data curation, Formal analysis, Investigation, Supervision, Validation, Writing—original draft, Writing— review & editing). P.P.B. (Conceptualization, Data interpretation, Funding acquisition, Supervision, Validation, Writing—original draft, Writing—review & editing).

## Supplementary data

Supplementary Data are available at journal’s site Online.

## Data availability

The mass spectrometry proteomics data have been deposited to the ProteomeXchange Consortium (http://proteomecentral.proteomexchange.org) via the PRIDE partner repository (43) with the dataset identifier PXD067195. Custom code for mass spectrometry analysis is available at https://github.com/ZApostolou/Krause_etal_2025.git.

