## Supplementary Data for "Intrinsic specificity of a ‘core’ Tip60 acetyltransferase complex in *Drosophila*"

### Supplementary Table 1

| Mass [m/z] | CS [z] | Polarity | (N)CE | (N)CE type | Comment |
|---|---|---|---|---|---|
| 508,6541 | 3 | Positive | 27 | NCE | H2A 1Ac |
| 507,6479 | 3 | Positive | 27 | NCE | H2A 2Ac |
| 506,6416 | 3 | Positive | 27 | NCE | H2A 3Ac |
| 505,6354 | 3 | Positive | 27 | NCE | H2A 4Ac |
| 670,7232 | 3 | Positive | 27 | NCE | H2AV 1Ac |
| 669,717 | 3 | Positive | 27 | NCE | H2AV 2Ac |
| 668,7107 | 3 | Positive | 27 | NCE | H2AV 3Ac |
| 667,7045 | 3 | Positive | 27 | NCE | H2AV 4Ac |
| 724,94279 | 2 | Positive | 27 | NCE | H4K5K8K12K16 1Ac |
| 723,43289 | 2 | Positive | 27 | NCE | H4K5K8K12K16 2Ac |
| 721,92206 | 2 | Positive | 27 | NCE | H4K5K8K12K16 3Ac |
| 685,733 | 3 | Positive | 27 | NCE | H2AV 1Ac + NTerm |
| 684,7267 | 3 | Positive | 27 | NCE | H2AV 2Ac + NTerm |
| 683,7205 | 3 | Positive | 27 | NCE | H2AV 3Ac + NTerm |
| 682,7143 | 3 | Positive | 27 | NCE | H2AV 4Ac + NTerm |

**Supplementary Table 1**. Inclusion list of precursor ions and related parameters used for targeted PRM analysis of H2A, H4 and H2A.V. CS, Charge state; (N)CE, Normalized Collision Energy

### Supplementary Figure 1

Formula for calculating the relative abundance of acetylated H2A peptides.

#### H2A Mono-Acetylation

y6 y4

$G\ K_5\ G\ |\ G\ K_8\ V\ |\ K_{10}\ |\ G\ K_{12}\ |\ A\ K_{14}\ S\ R$

b3 b6

$$\textbf{H2A\_noPTM} = \frac{\text{MS1\_G4R16\_noPTM\_precursor}}{\text{sum(MS1\_G4R16\_noPTM\_precursor, MS1\_G4R16\_1Ac\_precursor, MS1\_G4R16\_2Ac\_precursor)}}$$

- - - - - - - - - - - - - - - - - - - - - - - - - - - - - - - - - - - - - - - -

$$\textbf{H2A\_K14} = \frac{\text{MS1\_G4R16\_1Ac\_precursor}}{\text{sum(MS1\_G4R16\_noPTM\_precursor, MS1\_G4R16\_1Ac\_precursor, MS1\_G4R16\_2Ac\_precursor)}} * \frac{\text{MS2\_G2R16\_1Ac\_y4\_K14ac\_503\_y4}}{\text{sum(MS2\_G2R16\_1Ac\_y4\_K14ac\_503\_y4, MS2\_G2R16\_1Ac\_y4\_K14Noac\_506\_y4)}}$$

- - - - - - - - - - - - - - - - - - - - - - - - - - - - - - - - - - - - - - - -

$$\textbf{H2A\_K12} = \frac{\text{MS1\_G4R16\_1Ac\_precursor}}{\text{sum(MS1\_G4R16\_noPTM\_precursor, MS1\_G4R16\_1Ac\_precursor, MS1\_G4R16\_2Ac\_precursor)}} * \frac{\text{MS2\_G2R16\_1Ac\_y6\_K12ac\_733\_y6}}{\text{sum(MS2\_G2R16\_1Ac\_y6\_K12ac\_733\_y6, MS2\_G2R16\_1Ac\_y6\_K12Noac\_736\_y6)}} - \frac{\text{MS2\_G2R16\_1Ac\_y4\_K14ac\_503\_y4}}{\text{sum(MS2\_G2R16\_1Ac\_y4\_K14ac\_503\_y4, MS2\_G2R16\_1Ac\_y4\_K14Noac\_506\_y4)}}$$

- - - - - - - - - - - - - - - - - - - - - - - - - - - - - - - - - - - - - - - -

$$\textbf{H2A\_K10} = \frac{\text{MS1\_G4R16\_1Ac\_precursor}}{\text{sum(MS1\_G4R16\_noPTM\_precursor, MS1\_G4R16\_1Ac\_precursor, MS1\_G4R16\_2Ac\_precursor)}} * 1 - \frac{\text{MS2\_G2R16\_1Ac\_b6\_K8ac\_614\_b6}}{\text{sum(MS2\_G2R16\_1Ac\_b6\_K8ac\_614\_b6, MS2\_G2R16\_1Ac\_b6\_K8Noac\_617\_b6)}}$$

- - - - - - - - - - - - - - - - - - - - - - - - - - - - - - - - - - - - - - - -

$$\textbf{H2A\_K8} = \frac{\text{MS1\_G4R16\_1Ac\_precursor}}{\text{sum(MS1\_G4R16\_noPTM\_precursor, MS1\_G4R16\_1Ac\_precursor, MS1\_G4R16\_2Ac\_precursor)}} * \frac{\text{MS2\_G2R16\_1Ac\_b6\_K8ac\_614\_b6}}{\text{sum(MS2\_G2R16\_1Ac\_b6\_K8ac\_614\_b6, MS2\_G2R16\_1Ac\_b6\_K8Noac\_617\_b6)}} - \frac{\text{MS2\_G2R16\_1Ac\_b3\_K5ac\_285\_b3}}{\text{sum(MS2\_G2R16\_1Ac\_b3\_K5ac\_285\_b3, MS2\_G2R16\_1Ac\_b3\_K5Noac\_288\_b3)}}$$

- - - - - - - - - - - - - - - - - - - - - - - - - - - - - - - - - - - - - - - -

$$\textbf{H2A\_K5} = \frac{\text{MS1\_G4R16\_1Ac\_precursor}}{\text{sum(MS1\_G4R16\_noPTM\_precursor, MS1\_G4R16\_1Ac\_precursor, MS1\_G4R16\_2Ac\_precursor)}} * \frac{\text{MS2\_G2R16\_1Ac\_b3\_K5ac\_285\_b3}}{\text{sum(MS2\_G2R16\_1Ac\_b3\_K5ac\_285\_b3, MS2\_G2R16\_1Ac\_b3\_K5Noac\_288\_b3)}}$$

#### H2A Di-Acetylation

$G\ K_5\ G\ G\ K_8\ V\ |\ K_{10}\ G\ K_{12}\ A\ K_{14}\ S\ R$

b6

$$\textbf{H2A\_K5K8} = \frac{\text{MS1\_G4R16\_2Ac\_precursor}}{\text{sum(MS1\_G4R16\_noPTM\_precursor, MS1\_G4R16\_1Ac\_precursor, MS1\_G4R16\_2Ac\_precursor)}} * \frac{\text{MS2\_G2R16\_2Ac\_b6\_K5K8.2ac\_611\_b6}}{\text{sum(MS2\_G2R16\_2Ac\_b6\_K5K8.2ac\_611\_b6, MS2\_G2R16\_2Ac\_b6\_K5K8.1ac\_614\_b6, MS2\_G2R16\_2Ac\_b6\_K5K8.0ac\_617\_b6)}}$$

- - - - - - - - - - - - - - - - - - - - - - - - - - - - - - - - - - - - - - - -

$$\textbf{H2A\_K8K10 \_K5K10} = \frac{\text{MS1\_G4R16\_2Ac\_precursor}}{\text{sum(MS1\_G4R16\_noPTM\_precursor, MS1\_G4R16\_1Ac\_precursor, MS1\_G4R16\_2Ac\_precursor)}} * 1 - \frac{\text{MS2\_G2R16\_2Ac\_b6\_K5K8.2ac\_611\_b6}}{\text{sum(MS2\_G2R16\_2Ac\_b6\_K5K8.2ac\_611\_b6, MS2\_G2R16\_2Ac\_b6\_K5K8.1ac\_614\_b6, MS2\_G2R16\_2Ac\_b6\_K5K8.0ac\_617\_b6)}}$$

### Supplementary Figure 2

Formula for calculating the relative abundance of acetylated H2A.V peptides.

#### H2A.V Mono-Acetylation

y12 y8 y6

A G G $K_4$ | A G $K_7$ | D S G $K_{11}$ | A $K_{13}$ | A $K_{15}$ A V S R

b4

$$\text{H2A.V\_noPTM} = \frac{\text{MS1\_A1R19\_noPTM\_Ac\_NTerm\_precursor}}{\text{sum(MS1\_A1R19\_noPTM\_Ac\_NTerm\_precursor, MS1\_A1R19\_1Ac\_Ac\_NTerm\_precursor, MS1\_A1R19\_2Ac\_Ac\_NTerm\_precursor)}}$$

- - - - - - - - - - - - - - - - - - - - - - - - - - - - - - - - - - - - - - - - - -

$$\text{H2A.V\_K15} = \frac{\text{MS1\_A1R19\_1Ac\_Ac\_NTerm\_precursor}}{\text{sum(MS1\_A1R19\_noPTM\_Ac\_NTerm\_precursor, MS1\_A1R19\_1Ac\_Ac\_NTerm\_precursor, MS1\_A1R19\_2Ac\_Ac\_NTerm\_precursor)}} * \frac{\text{MS2\_A1R19\_1Ac\_y6\_K15ac\_673\_y6}}{\text{sum(MS2\_A1R19\_1Ac\_y6\_K15ac\_673\_y6, MS2\_A1R19\_1Ac\_y6\_K15Noac\_676\_y6)}}$$

- - - - - - - - - - - - - - - - - - - - - - - - - - - - - - - - - - - - - - - - - -

$$\text{H2A.V\_K13} = \frac{\text{MS1\_A1R19\_1Ac\_Ac\_NTerm\_precursor}}{\text{sum(MS1\_A1R19\_noPTM\_Ac\_NTerm\_precursor, MS1\_A1R19\_1Ac\_Ac\_NTerm\_precursor, MS1\_A1R19\_2Ac\_Ac\_NTerm\_precursor)}} * \left( \frac{\text{MS2\_A1R19\_1Ac\_y8\_K13ac\_917\_y8}}{\text{sum(MS2\_A1R19\_1Ac\_y8\_K13ac\_917\_y8, MS2\_A1R19\_1Ac\_y8\_K13Noac\_920\_y8)}} - \frac{\text{MS2\_A1R19\_1Ac\_y6\_K15ac\_673\_y6}}{\text{sum(MS2\_A1R19\_1Ac\_y6\_K15ac\_673\_y6, MS2\_A1R19\_1Ac\_y6\_K15Noac\_676\_y6)}} \right)$$

- - - - - - - - - - - - - - - - - - - - - - - - - - - - - - - - - - - - - - - - - -

$$\text{H2A.V\_K11} = \frac{\text{MS1\_A1R19\_1Ac\_Ac\_NTerm\_precursor}}{\text{sum(MS1\_A1R19\_noPTM\_Ac\_NTerm\_precursor, MS1\_A1R19\_1Ac\_Ac\_NTerm\_precursor, MS1\_A1R19\_2Ac\_Ac\_NTerm\_precursor)}} * \left( \frac{\text{MS2\_A1R19\_1Ac\_y12\_K11ac\_1349\_y12}}{\text{sum(MS2\_A1R19\_1Ac\_y12\_K11ac\_1349\_y12, MS2\_A1R19\_1Ac\_y12\_K11Noac\_1352\_y12)}} - \frac{\text{MS2\_A1R19\_1Ac\_y8\_K13ac\_917\_y8}}{\text{sum(MS2\_A1R19\_1Ac\_y8\_K13ac\_917\_y8, MS2\_A1R19\_1Ac\_y8\_K13Noac\_920\_y8)}} \right)$$

- - - - - - - - - - - - - - - - - - - - - - - - - - - - - - - - - - - - - - - - - -

$$\text{H2A.V\_K7} = \frac{\text{MS1\_A1R19\_1Ac\_Ac\_NTerm\_precursor}}{\text{sum(MS1\_A1R19\_noPTM\_Ac\_NTerm\_precursor, MS1\_A1R19\_1Ac\_Ac\_NTerm\_precursor, MS1\_A1R19\_2Ac\_Ac\_NTerm\_precursor)}} * \left( 1 - \frac{\text{MS2\_A1R19\_1Ac\_b4\_K4ac\_401\_b4}}{\text{sum(MS2\_A1R19\_1Ac\_b4\_K4ac\_401\_b4, MS2\_A1R19\_1Ac\_b4\_K4Noac\_404\_b4)}} \right)$$

- - - - - - - - - - - - - - - - - - - - - - - - - - - - - - - - - - - - - - - - - -

$$\text{H2A.V\_K4} = \frac{\text{MS1\_A1R19\_1Ac\_Ac\_NTerm\_precursor}}{\text{sum(MS1\_A1R19\_noPTM\_Ac\_NTerm\_precursor, MS1\_A1R19\_1Ac\_Ac\_NTerm\_precursor, MS1\_A1R19\_2Ac\_Ac\_NTerm\_precursor)}} * \frac{\text{MS2\_A1R19\_1Ac\_b4\_K4ac\_401\_b4}}{\text{sum(MS2\_A1R19\_1Ac\_b4\_K4ac\_401\_b4, MS2\_A1R19\_1Ac\_b4\_K4Noac\_404\_b4)}}$$

#### H2A.V Di-Acetylation

A G G $K_4$ A G $K_7$ D S | G $K_{11}$ A $K_{13}$ A $K_{15}$ A V S R

b9

$$\text{H2A.V\_K4K7ac} = \frac{\text{MS1\_A1R19\_2Ac\_Ac\_NTerm\_precursor}}{\text{sum(MS1\_A1R19\_noPTM\_Ac\_NTerm\_precursor, MS1\_A1R19\_1Ac\_Ac\_NTerm\_precursor, MS1\_A1R19\_2Ac\_Ac\_NTerm\_precursor)}} * \frac{\text{MS2\_A1R19\_2Ac\_b9\_K4K7.2ac\_901\_b9}}{\text{sum(MS2\_A1R19\_2Ac\_b9\_K4K7.2ac\_901\_b9, MS2\_A1R19\_2Ac\_b9\_K4K7.1ac\_904\_b9, MS2\_A1R19\_2Ac\_b9\_K4K7.0ac\_907\_b9)}}$$
